# Engineering Dual-Target Chimeric Lysins for Synergistic Eradication of *Porphyromonas gingivalis*

**DOI:** 10.64898/2026.04.27.720852

**Authors:** Fangfang Yao, Jiajun He, Fangyuan Chen, Tianshu Chu, Hongping Wei, Yuhong Li

**Affiliations:** The State Key Laboratory Breeding Base of Basic Science of Stomatology (Hubei-MOST) & Key Laboratory of Oral Biomedicine, Ministry of Education, School of Stomatology, Wuhan University, Wuhan 430079, China; Xiamen Key Laboratory of Stomatological Disease Diagnosis and Treatment, Stomatological Hospital of Xiamen Medical College, Xiamen, 361008, China; CAS-Key Laboratory of Synthetic Biology, CAS Center for Excellence in Molecular Plant Sciences, Institute of Plant Physiology and Ecology, Chinese Academy of Sciences, Shanghai 200032, China; University of Chinese Academy of Sciences, Beijing 100039, China; WHP Innovation Lab, Wuhan Institute of Virology, Chinese Academy of Sciences, Wuhan 430071, China

## Abstract

Periodontitis is a chronic inflammatory disease driven by complex subgingival multispecies biofilms, in which *Porphyromonas gingivalis* plays a central role in coordinating microbial interactions. Together with host-associated factors, these microbial communities create dual constraints that limit antimicrobial efficacy. In this study, we engineered a library of recombinant lysins by fusing membrane-destabilizing peptides to the periodontal pathogen–derived lysin LysPd078 and identified four optimized variants (PlyPd06, PlyPd19, PlyPd27, and PlyPd44) with enhanced salt tolerance, environmental stability, and bactericidal activity. In a clinically derived polymicrobial oral biofilm, PlyPd44 at only 25 μg/mL eradicated 96.6% of *P. gingivalis*. In a humanized oral microbiota mouse model of periodontitis, selected PlyPds significantly reduced inflammation and inhibited alveolar bone loss compared with LysPd078 and minocycline. Collectively, these results establish membrane-destabilizing peptide–enabled lysins as a promising platform for developing microenvironment-adapted precision antimicrobials for periodontitis.

## Introduction

Periodontitis is a chronic inflammatory disease driven by subgingival biofilms and represents a major cause of tooth loss in adults (Cui et al. 2023; Trindade et al. 2023). It is also closely associated with systemic conditions, including diabetes and cardiovascular diseases, making it an important public health concern (Peng et al. 2022; Villoria et al. 2024). Unlike classical infections caused by single pathogens, periodontitis is fundamentally a dysbiosis-driven disease characterized by disruption of highly organized multispecies microbial communities (Hajishengallis and Lamont 2021; Kang et al. 2021). Within this pathogenic network, *Porphyromonas gingivalis* plays a central role by reshaping community composition through extensive interspecies interactions and modulating host immune responses (Reyes 2021; Tang et al. 2024). Therefore, effective elimination of periodontal pathogens remains a key step in controlling disease progression. However, current clinical treatments, including scaling and root planing, show limited efficacy in removing pathogens embedded within subgingival biofilms (Herrera et al. 2022; Suvan et al. 2020).

A major challenge arises from the structural complexity of periodontal biofilms. These biofilms form protective matrices composed of extracellular polysaccharides, proteins, and nucleic acids that restrict antimicrobial diffusion and activity (Jakubovics et al. 2021; Kreth and Merritt 2023). Compared with planktonic cells, *P. gingivalis* within biofilms exhibits markedly reduced susceptibility to commonly used antimicrobials, including chlorhexidine, minocycline, and metronidazole, with tolerance differences reaching several hundred-to thousand-fold (Gerits et al. 2017). In addition to biofilm-mediated protection, host-associated microenvironments further compromise antimicrobial stability. Proteases such as gingipains degrade antimicrobial peptides including LL-37 (McCrudden et al. 2013), while salivary components and ionic conditions reduce the activity of endogenous peptides such as Histatin-5 (Dong et al. 2003). These findings suggest that periodontal lesions impose dual constraints consisting of biofilm-mediated and host-mediated barriers that limit antimicrobial efficacy under clinically relevant conditions (Jiao et al. 2019).

Despite these challenges, many antimicrobial candidates are still evaluated using single-species and simplified culture systems, which may systematically overestimate their translational potential (Soldati et al. 2023; Vyas et al. 2022). For example, the antimicrobial peptide Nal-P-113 exhibits a minimum inhibitory concentration of 20 μg/mL against *P. gingivalis* in vitro; however, in animal models, even 1280 μg/mL reduces bacterial counts by less than 1 log (Wang et al. 2015), and clinical studies have shown no significant improvement in clinical attachment loss following its application (Wang et al. 2018). These discrepancies highlight the importance of maintaining functional activity under host-associated conditions and underscore the need for experimental models that better replicate disease-relevant environments (Vyas et al. 2022).

Phage-derived lysins have emerged as promising antimicrobial agents due to their rapid bactericidal activity, ability to disrupt biofilms, and low risk of resistance development (McCallin et al. 2023). Several lysins targeting pathogens such as Staphylococcus aureus and Pseudomonas aeruginosa have progressed into clinical development, including Exebacase, which has reached phase III trials (Danis-Wlodarczyk et al. 2021; Ho et al. 2022). However, lysin development targeting periodontal pathogens is extremely rare (Kabwe et al. 2025). Recently, Xu and colleagues identified six candidate lysins targeting *P. gingivalis* from the human oral virome database (HOVD); nevertheless, even when used in combination at approximately 2.5 μM, these lysins achieved about 5% growth inhibition (Xu et al. 2025). In contrast, we previously identified a native lysin, LysPd078, derived from periodontal pathogens, which demonstrated substantially higher antibacterial activity under standard in vitro conditions, achieving a 99.99% reduction of *P. gingivalis* at 1.17 μM (Yao et al. 2026). Nevertheless, its activity was significantly reduced under simulated subgingival environments, suggesting that environmental robustness is a major limiting factor for lysin-based therapy.

The outer membrane of Gram-negative bacteria represents a key barrier restricting lysin access to the peptidoglycan layer, particularly under complex periodontal conditions where ionic strength, host proteins, and biofilm matrices further reduce lysin–bacterial interactions (Zheng and Zhang 2024). Previous studies have shown that introducing cationic or membrane-active peptide modules can enhance lysin interactions with bacterial outer membranes and improve antibacterial activity (Carratalá et al. 2023). However, whether this strategy can maintain effectiveness under periodontal-specific multispecies and host-associated environments remains insufficiently understood.

In this study, we established experimental systems that simulate clinically relevant periodontal lesion conditions to evaluate the performance of engineered lysins under multispecies biofilm and host-associated environments. We aimed to optimize the cooperative interaction between membrane-active modules and lysins to shift antibacterial performance from dependence on ideal in vitro conditions to stable functionality under dual biofilm and host-derived constraints. Using a multi-level evaluation framework, including patient-derived multispecies biofilm models and a humanized periodontal infection model, we systematically assessed the antibacterial efficacy and therapeutic potential of this engineered strategy under disease-relevant conditions.

## Material and Methods

Details on the materials and methods are provided in the supplementary appendix.

## Results

### Engineering and screening PlyPds with enhanced antibacterial robustness

A library of 44 peptide-fused LysPd078 variants was generated by fusing outer membrane–active peptides to the N terminus of LysPd078 (Fig. 1A and Supplementary Fig. 1). These peptides included lipopolysaccharide-binding peptides, antimicrobial peptides validated against periodontal pathogens, and peptides with reported activity against Gram-negative bacteria. Structural modeling suggested that N-terminal fusion did not alter the catalytic-site geometry or predicted peptidoglycan-binding interactions of LysPd078 (Fig. 1B-G and Supplementary Fig. 2). Of the 44 constructs, 17 were obtained in soluble form following expression in E. coli (Supplementary Fig. 3A). These soluble variants were screened against P. gingivalis W83 under buffer and high-salt conditions. Four variants, PlyPd06, PlyPd19, PlyPd27, and PlyPd44, showed the strongest activity and were selected for further analysis.

**Figure 1.**
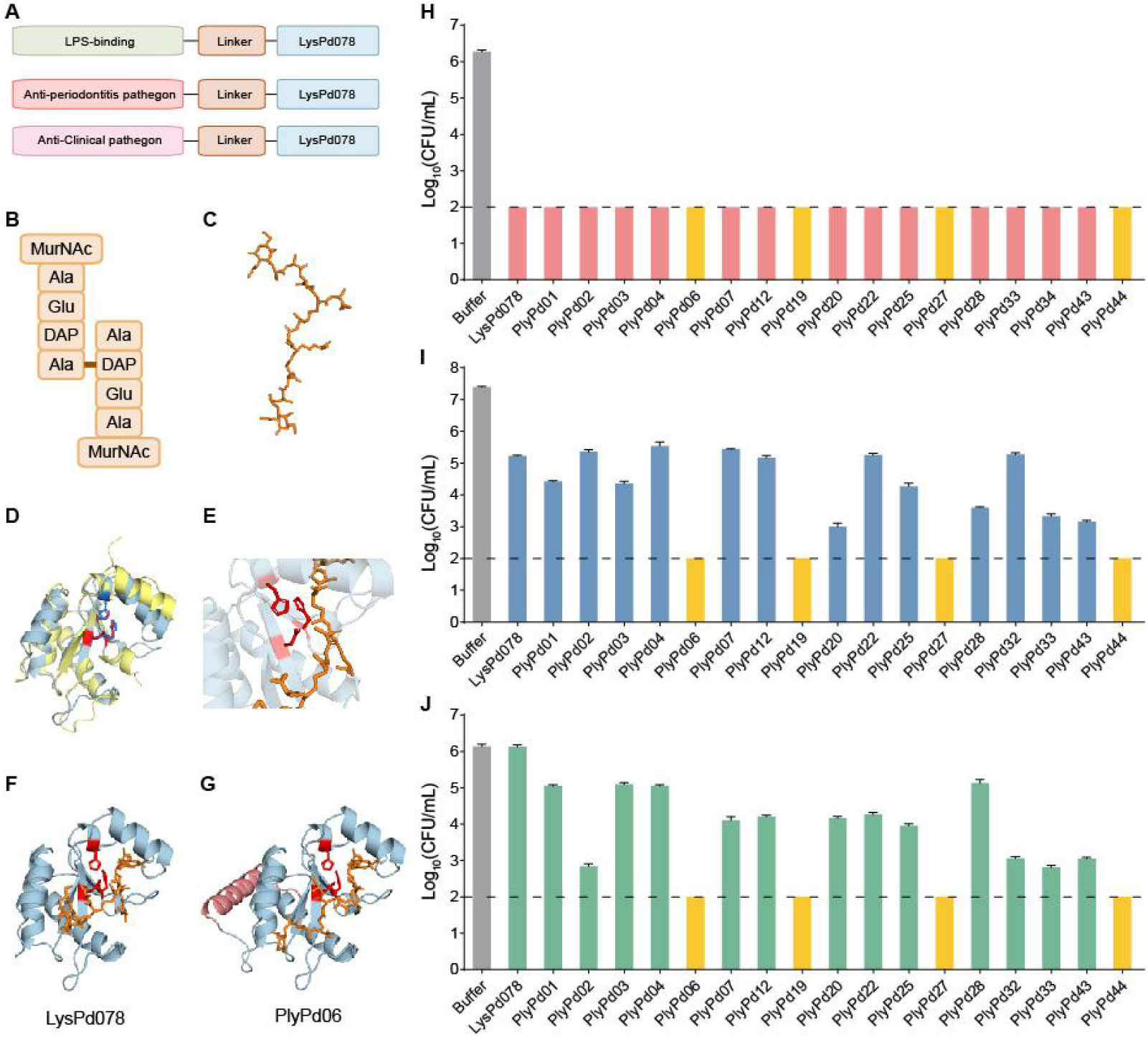
Structural characterization and antibacterial activity of engineered PlyPds. (A) Schematic representation of the designed PlyPd constructs. (B) Cartoon illustration of the peptidoglycan (PG) composition, including N-acetylmuramic acid (MurNAc), alanine (Ala), glutamic acid (Glu), and diaminopimelic acid (DAP). (C) Structural organization of PG. (D) Structural alignment of LysPd078 (blue) with its homolog LysECD7 (yellow). Catalytic residues of the peptidase active center are highlighted as stick models (red in LysPd078 and marine in LysECD7). (E) Close-up view of LysPd078 showing conserved catalytic residues (red sticks) and the modeled PG substrate (orange sticks). (F, G) Predicted binding conformations of LysPd078 (F) and engineered PlyPd06 (G) with PG. (H, I) Antibacterial activity against *P. gingivalis* W83 measured at bacterial concentrations of 10^6^ CFU/mL (H) and 10^7^ CFU/mL (I). (J) Antibacterial activity against *P. gingivalis* W83 in the presence of 150 mM NaCl. The dashed line indicates the detection limit.

Among engineered variants, 17 PlyPds were obtained in soluble form following E. coli expression (Supplementary Fig. 3A). Given the central role of P. gingivalis in periodontal dysbiosis, subsequent experiments were performed using P. gingivalis as the primary test organism. Peptidoglycan hydrolysis assays confirmed that catalytic activity was largely retained, with several constructs exhibiting enhanced hydrolytic efficiency compared with LysPd078 (Supplementary Fig. 3B; Fig. 1H). Antibacterial screening against P. gingivalis W83 showed that all soluble PlyPds maintained bactericidal activity comparable to or greater than the parental enzyme under buffer conditions. Notably, PlyPd06, PlyPd19, PlyPd27, and PlyPd44 reduced bacterial counts below the detection limit at high bacterial density (10^7^ CFU/mL), whereas LysPd078 achieved only limited reduction (Fig. 1I).

The advantage of engineered PlyPds became more pronounced under physiologically relevant salt conditions. Under150 mM NaCl, LysPd078 nearly lost antibacterial activity, whereas multiple PlyPds retained robust killing capacity, with the four leading variants reducing bacterial loads to undetectable levels (Fig. 1J). These data suggest that peptide fusion may improve salt tolerance. Accordingly, PlyPd06, PlyPd19, PlyPd27, and PlyPd44 were selected for further characterization.

### PlyPds maintain antibacterial activity under periodontal environmental and host-associated conditions

We next characterized the primary biochemical properties of the four leading PlyPds. All variants exhibited strong concentration-dependent bactericidal activity (Fig. 2A), with PlyPd06 and PlyPd27 reducing *P. gingivalis* viability below the detection limit at 12.5 μg/mL, indicating enhanced potency compared with LysPd078. Under high-salt conditions, 25–50 μg/mL PlyPds achieved complete bacterial clearance, whereas LysPd078 showed limited activity even at 400 μg/mL (Fig. 2B). Notably, whereas LysPd078 exhibited optimal activity primarily near neutral pH, leading PlyPds maintained strong antibacterial activity across a broader pH range (pH 6.0–10.0), indicating enhanced tolerance to pH fluctuations characteristic of inflammatory periodontal environments (Fig. 2C). Thermal stability was preserved following incubation from 20–100°C (Fig. 2D).

**Figure 2.**
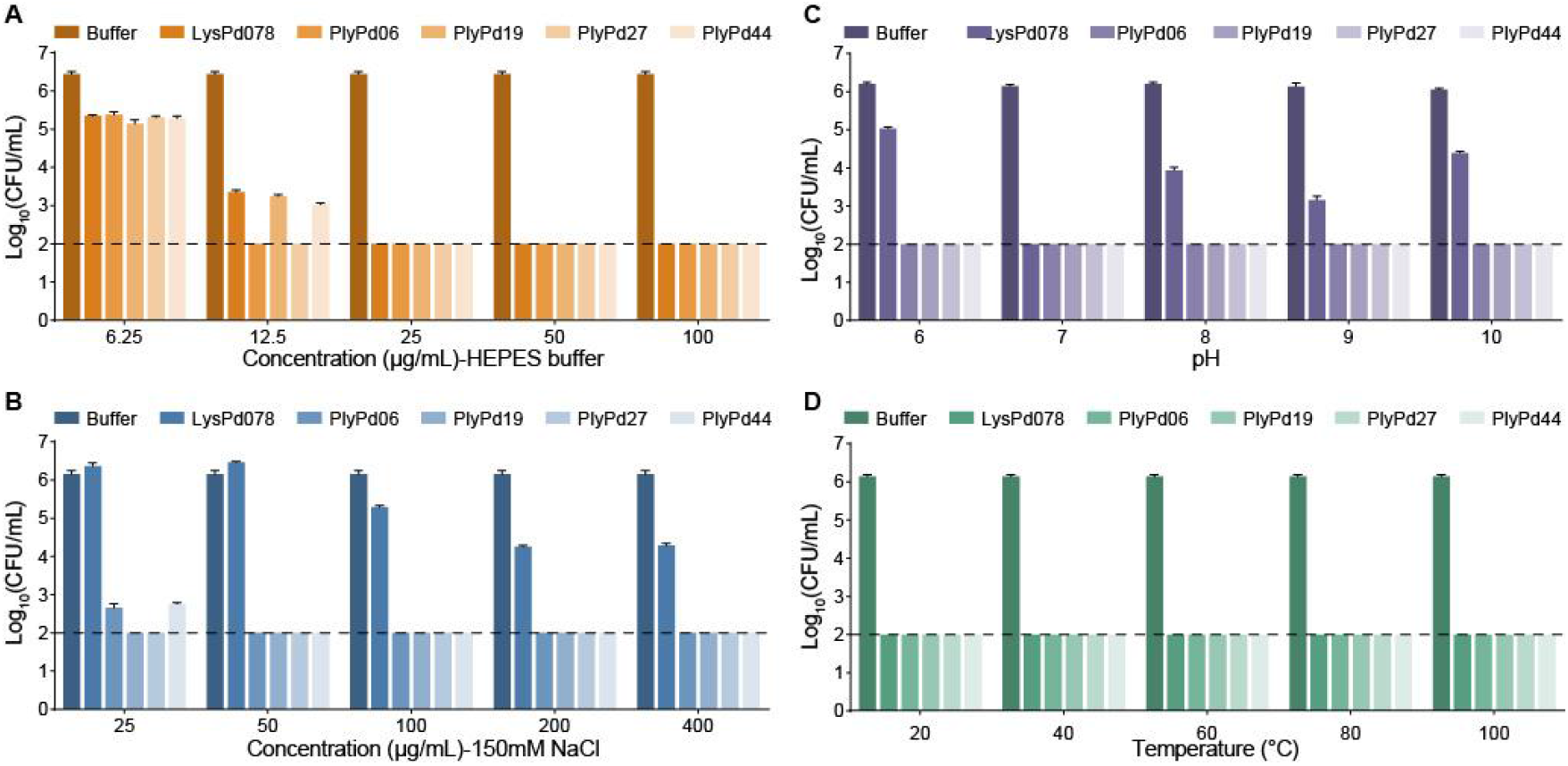
Bactericidal activity of LysPd078 and engineered PlyPds under physiologically relevant conditions. (A) Concentration-dependent bactericidal activity of LysPd078 and selected PlyPds in 10 mM HEPES buffer. (B) Bactericidal activity of LysPd078 and selected PlyPds in the presence of 150 mM NaCl. (C) Effect of pH on the bactericidal activity of LysPd078 and selected PlyPds at 100 μg/mL. (D) Effect of temperature on the bactericidal activity of LysPd078 and selected PlyPds at 100 μg/mL. The dashed line indicates the detection limit. Data are presented as mean ± SD.

Under host-associated periodontal conditions, we further evaluated the antibacterial activity of the selected PlyPds (Fig. 3). Simulated gingival crevicular fluid (sGCF) was prepared based on the concentrations of NaCl, KCl, CaCl_2_, and MgCl_2_ reported in the gingival crevicular fluid of patients with periodontitis, and was used to assess lysin activity under complex ionic conditions. Under these conditions, the selected PlyPds showed greater antibacterial activity than LysPd078. Specifically, 25 μg/mL PlyPd06, PlyPd27, and PlyPd44 reduced *P. gingivalis* counts below the detection limit, whereas LysPd078 required 100 μg/mL to achieve the same effect (Fig. 3A). In the presence of 10% human serum, the antibacterial activity of LysPd078 was markedly reduced, whereas PlyPd06 still decreased bacterial counts by approximately 3 log units at 50 μg/mL (Fig. 3B). After preincubation in human plasma for 10 h, the selected PlyPds largely retained their bactericidal activity, while LysPd078 showed a partial loss of activity (Fig. 3C). In addition, because proteolytic enzymes released by highly activated neutrophils in the gingival crevicular fluid of patients with periodontitis may affect lysin activity, we further examined antibacterial activity in the presence of elastase. Although 10 mU human elastase alone showed no antibacterial activity, the concentration of PlyPd06 required to reduce *P. gingivalis* W83 below the detection limit decreased to 6.25 μg/mL in its presence (Fig. 3D). Further analysis showed that the selected PlyPds retained strong antibacterial activity against most representative oral pathogens in sGCF, while exerting limited effects on representative commensal species (Fig. 3E). Together, these results indicate that engineered PlyPds exhibit improved functional robustness under host-associated periodontal conditions.

**Figure 3.**
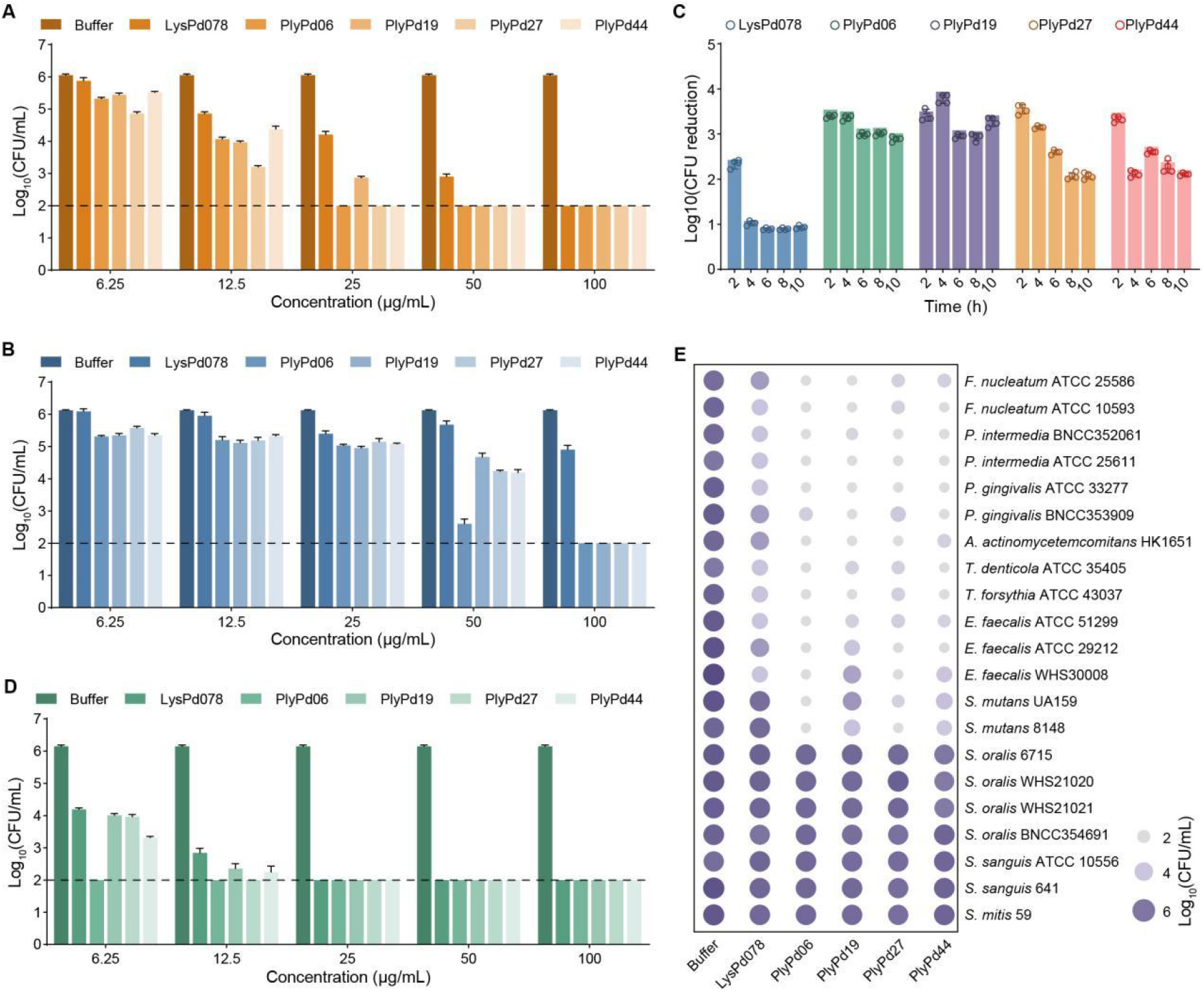
Biochemical characterization of LysPd078 and engineered PlyPds under conditions mimicking the periodontal microenvironment. (A,B) Effects of simulated gingival crevicular fluid (sGCF) (A) and serum (B) on the antibacterial activity of LysPd078 and selected PlyPds against *P. gingivalis* W83. (C) Antibacterial activity of selected PlyPds (25 μg/mL) against *P. gingivalis* W83 following plasma preincubation for the indicated times. (D) Antibacterial activity of LysPd078 and selected PlyPds against *P. gingivalis* W83 in the presence of elastase. (E) Antibacterial activity of LysPd078 and selected PlyPds (25 μg/mL) against representative oral pathogens and commensal bacteria under sGCF conditions. The dashed line indicates the detection limit. Data are presented as mean ± SD.

### Leading PlyPds exhibit potent antibacterial activity against *P. gingivalis* within clinically derived polymicrobial biofilms

To evaluate antibacterial efficacy under complex oral microbial conditions, a polymicrobial biofilm model was established using subgingival plaque samples obtained from patients with periodontitis, into which P. gingivalis was incorporated to recapitulate clinically relevant microbial community structures.

To enable accurate quantification of species-specific bacterial viability within biofilms, a PMAxx-assisted quantitative PCR (qPCR) assay was established and validated (Sereti et al. 2021). Serial dilution analyses of PMAxx-treated live cells and lysin-inactivated cells demonstrated a strong linear correlation between cycle threshold (Ct) values and bacterial concentrations, while PMAxx effectively suppressed amplification signals derived from dead cells, allowing reliable quantification of viable bacteria (Fig. 4A). This optimized assay was subsequently used to assess changes in P. gingivalis abundance following lysin treatment. Both LysPd078 and engineered PlyPds reduced *P. gingivalis* levels within polymicrobial biofilms in a dose-dependent manner (Fig. 4B–F). Notably, engineered PlyPds showed substantially greater activity than LysPd078 in patient-derived polymicrobial biofilms, with PlyPd44 reducing *P. gingivalis* abundance by 96.6% at 25 μg/mL compared with 32.0% for LysPd078. Total biofilm bacterial burden, measured by CFU enumeration, also decreased progressively with increasing treatment concentrations (Fig. 4G), indicating that engineered lysins retained robust antibacterial activity within complex microbial communities.

**Figure 4.**
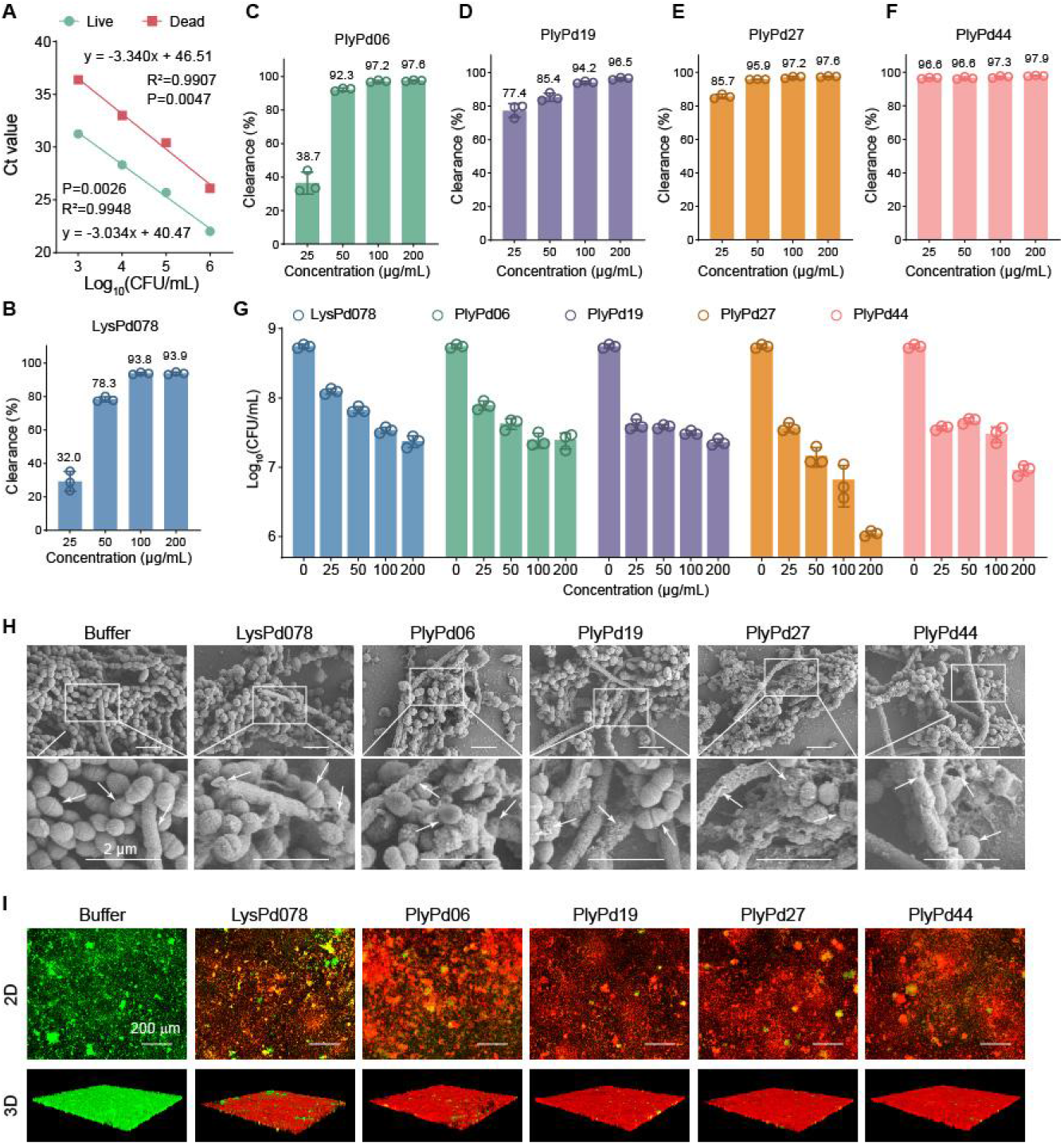
Antibacterial activity of LysPd078 and engineered PlyPds against *P. gingivalis* within clinically derived polymicrobial biofilms. (A) Linear regression analysis of Ct values obtained by PMAxx-assisted quantitative PCR (qPCR). Live and dead *P. gingivalis* cells were prepared by treatment with buffer or LysPd078, respectively, to establish calibration curves. (B–F) Bactericidal activity of LysPd078 (B), PlyPd06 (C), PlyPd19 (D), PlyPd27 (E), and PlyPd44 (F) against *P. gingivalis* embedded within polymicrobial biofilms derived from clinical plaque samples. Bacterial burden was quantified as the percentage of biofilm-associated cells killed by lysin treatment. (G) Total biofilm viability assessed by colony-forming unit (CFU) enumeration following treatment with LysPd078 and selected PlyPds. Data are presented as mean ± SD. (H) Representative live/dead fluorescence staining of treated biofilms. Green fluorescence indicates viable bacteria, and red fluorescence indicates dead bacteria. Scale bar, 100 μm. (I) Scanning electron microscopy (SEM) images showing morphological changes of biofilms after treatment with LysPd078 and selected PlyPds. Scale bar, 2 μm.

Scanning electron microscopy (SEM) was performed to investigate ultrastructural alterations. Untreated biofilms exhibited tightly packed microcolonies with intact cellular morphology, whereas lysin-treated samples showed pronounced structural disruption. Short rod-shaped and fusiform bacteria displayed surface perforations and fragmented debris (Fig. 4H), consistent with cell wall damage and bacterial lysis. Biofilm structural changes were further examined using live/dead fluorescence staining combined with confocal laser scanning microscopy (CLSM). Control biofilms displayed dense bacterial aggregates dominated by green fluorescence, indicating predominantly viable cells (Fig. 4I). In contrast, treatment with LysPd078 or engineered PlyPds resulted in marked increases in red fluorescence with localized orange signals, consistent with extensive bacterial death throughout the biofilm matrix (Fig. 4I). Collectively, species-specific quantification, community-level bacterial measurements, and multi-scale imaging analyses demonstrate that engineered PlyPds effectively reduce *P. gingivalis* abundance and disrupt biofilm architecture within clinically derived polymicrobial biofilms.

### Leading PlyPds alleviate periodontitis with superior efficacy to minocycline and favorable safety profiles

Prior to in vivo evaluation, cytocompatibility assays ddemonstrated that the four PlyPds exhibited excellent biocompatibility even at fivefold the therapeutic concentration, supporting their suitability for subsequent in vivo evaluation (Supplementary Fig. 4).

To evaluate in vivo antibacterial efficacy, we established a humanized murine periodontitis model by combining ligature placement with oral inoculation of *P. gingivalis* and subgingival plaque samples from patients with periodontitis, thereby partially recapitulating the complex microbial environment of human disease (Fig. 5A). Minocycline gel (Periocline), a clinically used periodontal therapeutic, was included as a positive control for comparison.

**Figure 5.**
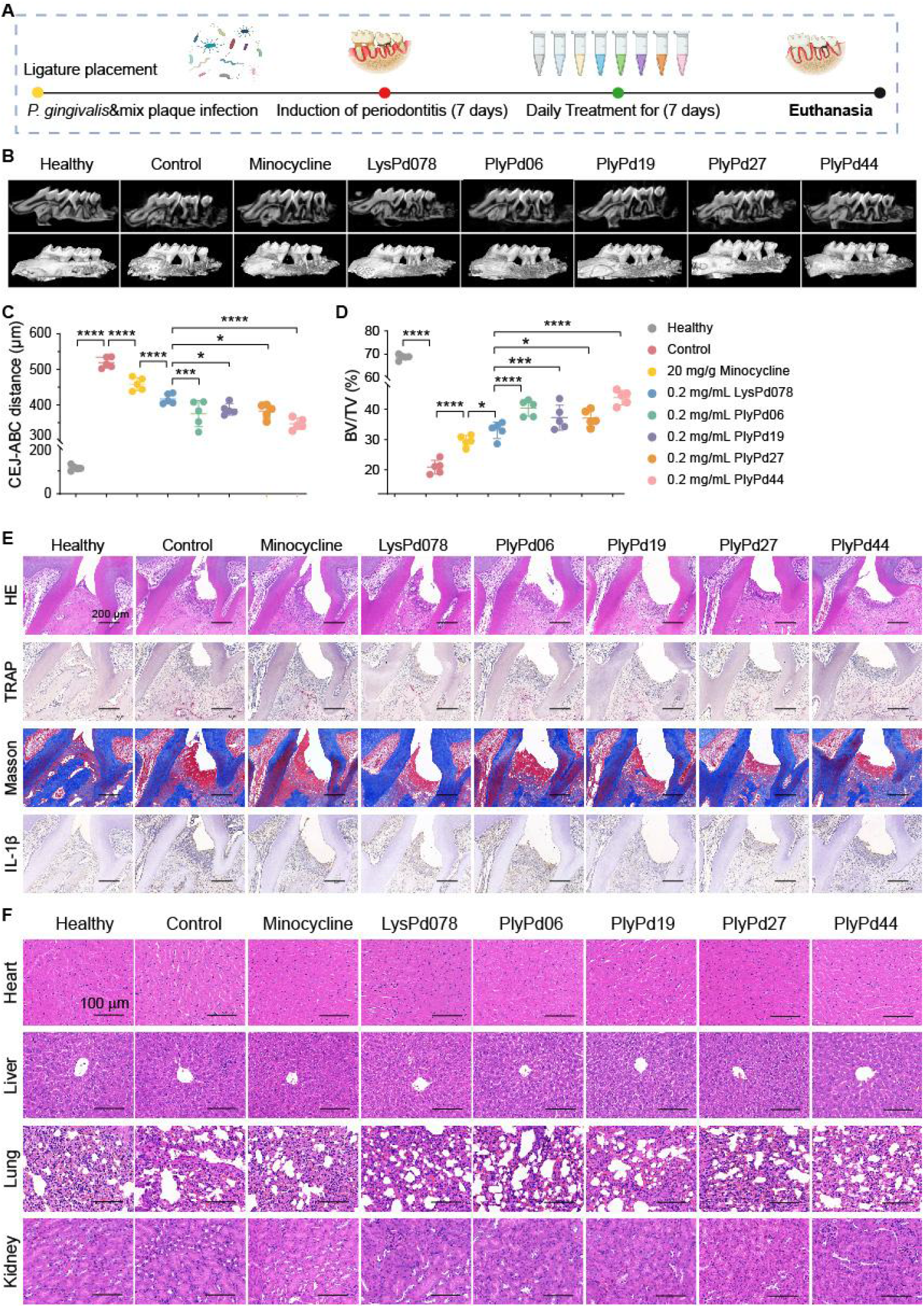
In vivo efficacy of LysPd078 and preferred PlyPds in a mouse periodontitis model infected with *P. gingivalis* W83 and subgingival plaque. (A) Schematic illustration of the periodontitis model, depicting the experimental design and treatment process. (B) Micro-CT reconstructed images of the right maxillary molar area from the indicated groups of mice. Both multiplanar reconstruction (Upper panels) and three-dimensional volume (Lower panels) are shown. (C) Quantification of the distance from the cementoenamel junction (CEJ) to the alveolar bone crest (ABC). (D) Ratio of bone volume (BV) to total volume (TV) of in the maxilla surrounding the second molar. (E) Histology staining images of periodontal tissue showing the pathogenic alteration. Scale bar, 100 μm. (F) Histological evaluation of representative organs (heart, liver, lung, kidney) stained with HE in periodontitis model mice. Scale bar, 100 µm. Statistical significance was calculated according to One Way ANOVA; n = 5/group, *p < 0.05, **p < 0.01, ***p < 0.001, ****p < 0.0001, ns, not significant. Each data point represents a biologically independent mouse.

Following successful model establishment, mice received daily treatments with minocycline, LysPd078, or engineered PlyPds (PlyPd06, PlyPd19, PlyPd27, and PlyPd44) for one week (Fig. 5B). Micro-computed tomography (micro-CT) analysis revealed severe alveolar bone destruction in untreated infected mice, characterized by pronounced furcation bone defects and extensive bone resorption extending toward the apical third of the roots. In contrast, treatment with LysPd078 or engineered PlyPds markedly reduced root exposure compared with both buffer and minocycline-treated groups (Fig. 5B).

Quantitative analyses further demonstrated significant increases in the CEJ–ABC distance and reductions in BV/TV values in the untreated group relative to healthy controls, confirming successful disease induction (Fig. 5C, D). Importantly, both LysPd078 and the engineered PlyPds exhibited more potent effects in alleviating alveolar bone loss compared to minocycline treatment. CEJ–ABC distance was significantly lower in the PlyPd44 group than in the minocycline group (P<0.001), with a corresponding increase in BV/TV.

Histological analyses of periodontal tissues surrounding the maxillary second molar further supported these findings. HE staining revealed extensive epithelial attachment loss and inflammatory cell infiltration in untreated mice, whereas lysin-treated groups displayed reduced inflammatory infiltration and improved tissue architecture (Fig. 5E). TRAP staining demonstrated decreased osteoclast activity following treatment, consistent with reduced bone resorption. Masson trichrome staining showed denser collagen fiber organization, indicating improved tissue integrity. Immunohistochemical analysis revealed markedly reduced IL-1β expression in lysin-treated groups compared with controls, onsistent with reduced inflammatory responses in lysin-treated mice (Fig. 5E).

Systemic safety was further evaluated throughout the treatment period. No significant differences in body weight were observed among treatment groups (Supplementary Fig. 5). Histological examination of major organs, including heart, liver, lung, and kidney, revealed no detectable pathological abnormalities (Fig. 5F), indicating favorable in vivo biocompatibility.

These findings show that engineered PlyPds reduced periodontal inflammation, suppressed osteoclast activation, and limited alveolar bone loss in the humanized polymicrobial periodontitis model without detectable systemic toxicity. Among the tested variants, PlyPd44 showed the strongest therapeutic effect.

### Discussion

An ideal antimicrobial strategy for periodontitis should maintain functional activity within the complex periodontal microenvironment, selectively eliminate key pathogens embedded in subgingival biofilms, and ultimately attenuate inflammation and alveolar bone loss (Siddiqui et al. 2023). However, periodontal lesions represent not merely sites of bacterial accumulation but dynamic ecosystems shaped by structured multispecies biofilms, host inflammatory responses, and diverse physicochemical factors (Hajishengallis and Lamont 2021). This environment promotes persistent pathogen enrichment and biofilm maturation while simultaneously reducing antimicrobial performance through elevated ionic strength, host-derived fluids, and proteolytic enzymes (Jakubovics et al. 2021). Within this context, we engineered a library of peptide-fused lysins based on the previously identified periodontal lysin LysPd078 and systematically evaluated their antibacterial and therapeutic performance under disease-relevant conditions. Our findings demonstrate that membrane-active peptide fusion substantially improves the functional robustness of lysins in complex periodontal environments.

Notably, the key improvement introduced by membrane-active peptide fusion extended beyond enhanced antibacterial activity under standard laboratory conditions (Carratalá et al. 2023). Instead, the primary benefit was a marked increase in functional robustness under complex environmental stressors relevant to periodontal lesions. The optimized PlyPds maintained significantly higher antibacterial activity than the parental lysin across elevated salt concentrations, broad pH ranges, and in the presence of simulated gingival crevicular fluid, serum, plasma, and neutrophil elastase. These findings indicate that peptide fusion enhances tolerance to both physicochemical and host-derived inhibitory factors that are known to compromise antimicrobial efficacy in *vivo*. Importantly, these results shift the focus from maximizing antibacterial potency under simplified conditions to optimizing sustained functionality under disease-relevant environments.

The enhanced environmental robustness observed in vitro translated into improved performance in clinically relevant polymicrobial biofilms. Unlike conventional assessments that rely solely on total CFU or standard qPCR measurements, our study employed a multilayered evaluation strategy combining PMAxx-qPCR, CFU enumeration, and multiscale imaging to characterize antibacterial efficacy at species-specific, community-wide, and structural levels. Within patient-derived multispecies biofilms, engineered PlyPds significantly reduced the abundance of *P. gingivalis*, accompanied by reductions in total viable biomass and disruption of biofilm architecture. Among the engineered variants, PlyPd44 achieved a 96.6% reduction of *P. gingivalis* at 25 μg/mL, markedly outperforming the parental lysin under identical conditions. These observations indicate that improvements in environmental stability are directly translated into enhanced pathogen clearance within complex microbial communities.

More importantly, the functional advantages observed under complex in vitro conditions translated into measurable therapeutic benefits in vivo. In the humanized polymicrobial periodontitis model established using ligature placement, *P. gingivalis* inoculation, and patient-derived subgingival microbiota, both LysPd078 and engineered PlyPds attenuated alveolar bone loss and inflammatory tissue damage. Micro-CT and histological analyses consistently demonstrated reductions in CEJ–ABC distance and improvements in BV/TV, accompanied by decreased inflammatory cell infiltration, reduced TRAP-positive osteoclast activity, and lower IL-1β expression. Notably, compared with locally administered minocycline, both the parental lysin and optimized PlyPds exhibited superior bone-protective effects, with PlyPd44 showing the most pronounced therapeutic efficacy. These findings suggest that membrane-active peptide fusion not only enhances antibacterial activity but also improves therapeutic performance in clinically relevant disease models. In addition, no significant changes in body weight or major organ histology were observed during short-term treatment, indicating favorable in vivo tolerability under the tested conditions.

Despite these encouraging findings, several limitations should be acknowledged. In the humanized polymicrobial model, engineered PlyPds significantly reduced pathogen burden and improved bone destruction and inflammatory phenotypes; however, inflammatory infiltration and osteoclast activation were not completely eliminated. Compared with simplified infection systems used in our previous studies, the inclusion of patient-derived subgingival microbiota resulted in more severe periodontal destruction, highlighting the increased biological complexity associated with multispecies interactions. This observation suggests that the structural organization and ecological stability of complex subgingival biofilms may continue to limit complete microbial eradication, even when pathogen loads are substantially reduced. Consequently, increasing ecological complexity may prolong immune activation and constrain full tissue recovery in advanced disease settings.

These findings provide important directions for future optimization. Further enhancement of lysin penetration and clearance depth within mature polymicrobial biofilms may help overcome the structural protection conferred by complex communities (Kwon et al. 2021; Suvan et al. 2020). In addition, optimizing local delivery and retention strategies may prolong therapeutic exposure within periodontal lesions and improve treatment durability (Wei et al. 2021; Zhao et al. 2020). Finally, integrating antimicrobial enhancement with host-directed anti-inflammatory modulation may further improve outcomes in environments characterized by persistent immune activation (Das et al. 2024; Petruk et al. 2026). Advances in structure-guided or artificial intelligence–assisted protein engineering may facilitate improvements in catalytic efficiency, substrate accessibility, and environmental stability (Hou et al. 2025; Stock and Gorochowski 2024; Zheng et al. 2025), while large-animal models that more closely resemble clinical conditions will be essential for evaluating pharmacokinetics, long-term safety, and translational potential (Mukherjee et al. 2022).

In summary, our study demonstrates that membrane-active peptide fusion substantially enhances the functional robustness of lysins under complex periodontal lesion environments and enables improved antibacterial and therapeutic performance in both patient-derived polymicrobial biofilms and humanized periodontitis models. These findings highlight that, for diseases driven by structured biofilms and host-associated constraints, improving antimicrobial activity under ideal laboratory conditions alone is insufficient to ensure therapeutic success. Instead, strengthening sustained functional performance under disease-relevant environments represents a critical design principle. Accordingly, environmentally robust engineered lysins may provide a promising strategy for precision antimicrobial therapy targeting periodontal diseases.

## Acknowledgments

This work was supported by the National Natural Science Foundation of China (No. 81870756). We thank Dr. Xuefang An, Dr. Tao Zhand, and Dr. Li, Li at the Center for Experimental Animals, Wuhan Institute of Virology for their assistance in animal experiments. We are grateful to Dr. Ding Gao, and Dr.Bichao Xu at the Center for Instrumental Analysis and Metrology, Wuhan Institute of Virology, CAS for technical assistance in transmission electron microscopy, scanning electron microscopy, and Confocal laser scanning microscopy.

## Author information

These authors contributed equally: Fangfang Yao, Jiajun He.

## Contributions

F.Y., J.H., H.W., and Y.L. designed the study. J.H. performed the bioinformatics work. F.Y., F.C., and T.C., carried out the experiments. F.Y., J.H., H.W., and Y.L. analyzed the data. F.Y., J.H. drafted the manuscript. F.Y., J.H., H.W., and Y.L. edited the manuscript. All authors gave their final approval and agree to be accountable for all aspects of the work.

## Competing interests

The authors declare no competing financial interests.

## Notes

### Competing Interest Statement

The authors have declared no competing interest.

