## Supplementary for "Engineering Dual-Target Chimeric Lysins for Synergistic Eradication of *Porphyromonas gingivalis*"

### Materials and methods

#### Bacterial strains and culture conditions

Bacterial strains, media, sources, and culture conditions used in this study are summarized in Table 1. Depending on experimental requirements, bacteria were cultured under aerobic, microaerophilic (5% $CO_2$), or anaerobic conditions using an AnaeroPack system (Mitsubishi).

**Table 1. Bacterial strains and growth conditions.**

| Species | Strain | Medium[a] | Condition[b] | Source[c] |
|---|---|---|---|---|
| Oral pathogens | | | | |
| *Aggregatibacter actinomycetemcomitans* | HK1651 | 2 | 3 | 2 |
| *Enterococcus faecalis* | ATCC 29212 | 1 | 3 | 2 |
| | ATCC 51299 | 1 | 3 | 2 |
| | WHS30008 | 1 | 3 | 1 |
| *Fusobacterium nucleatum* | ATCC 10953 | 2 | 4 | 2 |
| | ATCC 25586 | 2 | 4 | 2 |
| *Porphyromonas gingivalis* | ATCC 33277 | 2 | 4 | 2 |
| | BNCC353909 | 2 | 4 | 2 |
| | W83 | 2 | 4 | 2 |
| *Prevotella intermedia* | ATCC 25611 | 2 | 4 | 2 |
| | BNCC352061 | 2 | 4 | 2 |
| *Streptococcus mutans* | UA159 | 1 | 3 | 2 |
| | 8148 | 1 | 3 | 2 |
| *Treponema denticola* | ATCC 35405 | 3 | 4 | 3 |
| *Tannerella forsythia* | ATCC 43037 | 4 | 4 | 4 |
| Oral commensals | | | | |
| *Streptococcus mitis* | 59 | 1 | 3 | 2 |
| *Streptococcus oralis* | BNCC354691 | 1 | 3 | 2 |

| | | | | |
|---|---|---|---|---|
| | 6715 | 1 | 3 | 2 |
| | WHS21020 | 1 | 3 | 1 |
| | WHS21021 | 1 | 3 | 1 |
| *Streptococcus sanguis* | 641 | 1 | 3 | 2 |
| | ATCC 10556 | 1 | 3 | 2 |
| Other | | | | |
| *Escherichia coli* | BL21 (DE3) | 5 | 1 | 1 |

[a]Media: 1. Tryptic Soy Broth. 2.Tryptic Soy Broth supplemented with 5 mg/mL yeast extract, 0.5 mg/mL L-cysteine hydrochloride, 5 μg/mL hemin and 1 μg/mL menadione. 3. Fastidious Anaerobe Broth (Solarbio, China). 4. Fastidious Anaerobe Broth supplemented with 1 μg/mL MurNAc. 5. Lysogeny broth (LB). 6. Tryptic Soy Broth with 5% FBS. 7. Lysogeny broth (LB) with 0.1% borate. 8. Casamino acid-Peptone-Glucose broth. 9. de Man, Rogosa and Sharpe Broth. The agar plate of media 2, 3, 4, 6 were supplemented with 5% defibrinated sheep blood.

[b]Condition: 1. Aerobic cultivation, 37°C with aeration (200 rpm). 2. Aerobic cultivation, 28°C with aeration (200 rpm). 3. Microaerophilic cultivation, 37°C under 5% $CO_2$, 4. Anaerobic cultivation, 37°C with AnaeroPack™ (Mitsubishi, Japan).

[c]Source: 1. Key Laboratory of Special Pathogens and Biosafety, Wuhan Institute of Virology, China. 2. Stomatological Hospital of Wuhan University, China. 3. Oral Microbiome Bank of China. 4. Shanghai Ninth People's Hospital of Shanghai Jiao Tong University, China.

**Construction of expression plasmids for chimeric lysins**

All outer membrane-destabilizing peptides (IMPs) and engineered chimeric lysins included in this study are summarized in Table 2. LysPd078, a lysin active against periodontitis-associated pathogens, was previously generated in our laboratory. To construct the chimeric lysins, homologous overlap sequences were introduced into the peptide-coding fragments and the pET-28b (+) vector. The DNA fragments encoding each peptide or lysin variant were amplified by PCR using specific primers, purified, and quantified, and then cloned into the linearized vector using the ClonExpress Ultra One Step Cloning Kit. Through this strategy, the selected peptides were fused to the N terminus of LysPd078, yielding the recombinant engineered lysins.

**Table 2. Overview of engineered LysPd078.**

| Artilysin | IMP | IMP sequence | PMCID |
|---|---|---|---|
| PlyPd01 | Li5-001 | KNYSSSISSIHAC | 20816904 |
| PlyPd02 | LPS binding motif | GWKRKRFG | PMC4023739 |
| PlyPd03 | $\beta_2$GPI-V | AFWKTDA | 21947351 |
| PlyPd04 | Lfcin (28-34) | RKVRGPP | 27242337 |
| PlyPd05 | Peptide 2 | LVGKLLKGAVGDVCGLLPIC | 33689880 |
| PlyPd06 | LL-37 (7-27) | RKSKEKIGKEFKRIVQRIKDF | 20338846 |
| PlyPd07 | DP7 | VQWRIRVAVIRK | 35567537 |
| PlyPd08 | Pep-7 | RPHGAGEGIDRVPAGPSPSEVGLAIPSGK | 26041027 |
| PlyPd09 | SMAP29 | RGLRRLGRKIAHGVKKYGPTVLRIIRIAG | PMC356847 |
| PlyPd10 | LPcin-YK3 | NKVKEWIKYLKSLFS | 30021422 |
| PlyPd11 | Tachyplesin-1 [I11A] | KWCFRVCYRGACYRRCR | 28960954 |
| PlyPd12 | SynSaf-P56 | AGKEKIRKKLKNEIKKKWRKAVIAW | MC7296547 |
| PlyPd13 | HX-12C | FFRKVLKLIRKIWR | 29663932 |
| PlyPd14 | ApoE23 | LRKLRKRLVRLASHLRKLRKRLL | 24243597 |
| PlyPd15 | α4-Short | LKKWWKTSKGLLGGLLGKVTSVIK | PMC6456746 |
| PlyPd16 | Pyrrhocoricin [L7R, R14K] | VDKGSYRPRPTPPKPIYNRN | PMC6726534 |
| PlyPd17 | Thanatin | GSKKPVPIIYCNRRTGKCQRM | PMC4914623 |

| | | | |
|---|---|---|---|
| PlyPd18 | Microcin J25 | GGAGHVPEYFVGIGTPISFYG | PMC8894198 |
| PlyPd19 | Histatin5 | DSHAKRHHGYKRKFHEKHHSHRGY | 37395749 |
| PlyPd20 | E7K_I9K_IsCT | ILGKIWKGKKSLF | 25773522 |
| PlyPd21 | Ci-MAM-A24 | WRSLGRTLLRLSHALKPLARRSGW | 18598239 |
| PlyPd22 | SSL-25 | SSLLEKGLDGAKKAVGGLGKLGKDA | 19201578 |
| PlyPd23 | EC1-17KV | GWWRRTVKKVRNAVRKV | PMC7218270 |
| PlyPd24 | Kunitzin-RE W8 | AARIILRWRFR | 30958004 |
| PlyPd25 | GL13K | GKIIKLKASLKLL | MC3356437 |
| PlyPd26 | Cathelicidin-BF | KFFRKLKKSVKKRAKEFFKKPRVIGV SIPF | 25807257 |
| PlyPd27 | Ω76 | FLKAIKKFGKEFKKIGAKLK | PMC6656545 |
| PlyPd28 | PaDBS1R6 | PMARNKKLLKKLRLKIAFK | 30926365 |
| PlyPd29 | R7I | IRPIIRPIIRPIIRPIIRPIIRPIIRPI | 30742437 |
| PlyPd30 | hLF1-11 | GRRRRSVQWCA | 21827807 |
| PlyPd31 | KSL-W | KKVVFWVKFK | PMC4608580 |
| PlyPd32 | BP100 | KKLFKKILKYL | 23246973 |
| PlyPd33 | CecropinB | KWKVFKKIEKMGRNIRNGIVKAGPAI AVLGEAKAL | PMC1072432 7 |
| PlyPd34 | CecropinA | KWKLFKKIEKVGQNIRDGIIKAGPAV AVVGQATQIAK | PMC9610619 |
| PlyPd35 | CecropinA-8 | RWKIFKKI | 21593770 |
| PlyPd36 | Sarcotoxin IA | GWLKKIGKKIERVGQHTRDATIQGLGI AQQAANVAATARG | 3182836 |
| PlyPd37 | Chrysophsin-1 | FFGWLIKGAIHAGKAIHGLIHRRRH | PMC5469811 |
| PlyPd38 | TB_KKG6A | KKLLPIVANLLKSLL | 23403136 |
| PlyPd39 | Melittin | GIGAVLKVLTTGLPALISWIKRKRQQ | PMC3761581 |
| PlyPd40 | Grammistin Gs B | IGGIISFFKRLF | 15777955 |

| | | | |
|---|---|---|---|
| PlyPd41 | Buforin-2 (10-21) | FPVGRVHRLLRK | PMC26932 |
| PlyPd42 | Gaegurin 5 (1-13)[G3K] | FLKALFKVASKVL | 14739294 |
| PlyPd43 | Buforin-2 (8-21) | LQFPVGRVHRLLRK | PMC26932 |
| PlyPd44 | Thanatin 3 | GSKKPVPIIYCNRRKCQRM | 33546369 |

**Modeling of lysin-peptidoglycan complexes**

The canonical peptidoglycan structure of Gram-negative bacteria was drawn in ChemDraw. Chai-1 was used to predict the structures of lysin-peptidoglycan complexes and to analyze their interactions. Structural visualization and rendering were performed in PyMOL.

**Recombinant expression and purification of lysins**

Recombinant lysins were expressed in Escherichia coli BL21 cells as previously described with minor modifications. Briefly, transformed cells were induced with 0.2 mmol/L IPTG at 16 °C for 14 h. Cells were then harvested and disrupted by sonication on ice. Recombinant proteins were purified using HisTrap FF columns (GE Healthcare, Chicago, IL, USA) according to the manufacturer's instructions. Proteins were washed with 40 mM imidazole and eluted with 250 mM imidazole, followed by dialysis against 10 mM HEPES buffer (pH 7.4). Purified proteins were stored at 4 °C until use. Protein concentrations were determined using a Pierce BCA Protein Assay Kit (Thermo Scientific, Waltham, MA, USA).

**Peptidoglycan hydrolysis assay**

Briefly, bacteria were collected by centrifugation at 8,000 rpm for 5 min, boiled in 5% SDS with vigorous agitation, incubated overnight at room temperature, washed, and resuspended in ultrapure water, and stored at 4°C until use. LysPd078 or PlyPds were then added to the crude peptidoglycan suspension at a final concentration of 100 μg/mL and incubated at 37 °C for 1 h. OD600 was measured using a Synergy H1 microplate reader, with 10 mM HEPES buffer as the negative control. Relative peptidoglycan hydrolytic activity was calculated as: Ratio = (ODbuffer-ODnc) / (ODsample -ODnc), where ODnc is the OD600 of the negative control, ODbuffer is the OD600 of the buffer-treated sample, and ODsample is the OD600 of the lysin-treated sample.

**Bactericidal activity assay**

Bactericidal activity was assessed by colony-forming unit (CFU) counting. Log-phase Porphyromonas gingivalis W83 cells were harvested, washed, and resuspended in 10 mM HEPES buffer (pH 7.4) to approximately $10^6$ CFU/mL unless otherwise specified. Equal volumes of bacterial suspension and lysin solution or buffer control were mixed in 96-well plates and incubated at 37 °C for 1 h. Surviving bacteria were quantified by serial dilution and spot plating on blood agar plates. Samples were diluted immediately after treatment to minimize carryover killing during plating.

To evaluate the activities of different PlyPds against *P. gingivalis* W83, assays were performed using bacterial suspensions at approximately $10^6$ or $10^7$ CFU/mL in HEPES buffer, or $10^6$ CFU/mL in HEPES supplemented with 300 mM NaCl, with lysins applied at a final concentration of 100 μg/mL for 1 h at 37 °C. For comparative characterization of LysPd078 and PlyPds, bactericidal activity was further examined under different lysin concentrations, salt conditions, temperatures, pH values, and periodontal microenvironment-mimicking conditions, including simulated gingival crevicular fluid (sGCF), 10% serum, elastase, and plasma preincubation. Specifically, lysin concentrations ranged from 0 to 100 μg/mL in HEPES buffer and from 0 to 400 μg/mL in HEPES containing 150 mM NaCl. Thermal stability was assessed after preincubation at 20-100 °C for 1 h. pH-dependent activity was examined in sodium acetate buffer (pH 6.0), HEPES buffer (pH 7.0), Tris-HCl buffer (pH 8.0-9.0), and glycine-NaOH buffer (pH 10.0). For sGCF, serum, and elastase assays, lysins were tested at final concentrations of 0-100 μg/mL, with elastase used at a final concentration of 10 mU. For plasma tolerance assays, lysins were preincubated at 25 μg/mL for the indicated times before bactericidal testing.

The antibacterial spectrum of LysPds was further assessed against different oral pathogens and commensal bacteria.

**Mixed plaque biofilm model and PMAxx-qPCR analysis**

This study was approved by the Medical Ethics Committee of the School of Stomatology, Wuhan University (2024LUNSHENA32), and written informed consent was obtained from all participants before enrollment. Periodontal examination was performed before sampling by a trained examiner. Individuals were included if they showed gingival bleeding within 15 s after probing, had at least one site with a probing pocket depth > 5 mm, and had at least one site with clinical attachment loss > 4 mm. Individuals who had used antibiotics or probiotics, smoked, or received periodontal treatment within the previous 6 months were excluded. Subgingival plaque (SP) samples were pooled, mixed with an equal volume of 50% glycerol, aliquoted into sterile cryovials, and stored at −80 °C until use

To establish mixed plaque biofilms, *P. gingivalis* W83 was mixed 1:1 with resuspended SP samples and adjusted to approximately $10^7$ CFU/mL. The mixture was incubated anaerobically at 37 °C for 72 h. Biofilms were then washed with 10 mM HEPES buffer to remove non-adherent cells and treated with buffer alone or LysPd078/PlyPds at final concentrations of 25, 50, 100, or 200 μg/mL for 1 h at 37 °C. To quantify viable *P. gingivalis* in treated biofilms, two control groups were prepared: buffer-treated *P. gingivalis* as the live-cell control and *P. gingivalis* treated with 100 μg/mL LysPd078 as the dead-cell control. Viability was confirmed by CFU counting. After treatment, biofilms were thoroughly resuspended by repeated pipetting and vortexing. Samples were then treated with 100 μM PMAxx for 10 min, followed by light exposure for 10 min, to suppress amplification of DNA derived from membrane-compromised dead cells. Genomic DNA was extracted using a bacterial genomic DNA extraction kit. Viable *P. gingivalis* was quantified by qPCR targeting the waaA gene using the following primer/probe set: forward, 5′-TGGTTTCATGCAGCTTCTTT-3′; reverse, 5′-TCGGCACCTTCGTAATTCTT-3′; probe, 5′-FAM-CGTACCTCATATCCCGAGGGGCTG-BHQ-3′. Each 20 μL reaction contained 5 μL template DNA, 10 μL 2× reaction mix, 0.4 μM of each primer, 0.2 μM probe, and DNase-free water. Amplification was performed at 95 °C for 5 min, followed by 40 cycles of 95 °C for 10 s and 60 °C for 30 s. Standard curves generated from the live- and dead-cell controls were used to determine the abundance of viable *P. gingivalis* in mixed biofilms.

**CFU enumeration of mixed plaque biofilms**

Mixed plaque biofilms without PMAxx treatment were thoroughly resuspended by repeated pipetting and vortexing to disperse adherent bacteria. Serial dilutions were plated separately under aerobic and anaerobic conditions, and the CFU counts obtained from both conditions were pooled to quantify the total viable bacterial load remaining after treatment.

**Confocal laser scanning microscopy**

Mixed plaque biofilms were formed in 24-well plates containing coverslips as described previous. Biofilms were treated with buffer, 200 μg/mL LysPd078, or 200 μg/mL PlyPds for 1 h at 37 °C, and then stained using the Live & Dead Bacterial Staining Kit at room temperature. Images were acquired by confocal laser scanning microscopy (UltraVIEW VoX; PerkinElmer). Live bacteria are shown in green and dead bacteria in red.

**Scanning electron microscopy**

Mixed plaque biofilms grown on coverslips in 24-well plates were treated with buffer, 200 μg/mL LysPd078, or 200 μg/mL PlyPds for 1 h at 37 °C. Samples were fixed overnight at 4 °C in 2.5% glutaraldehyde, post-fixed with osmium tetroxide, and dehydrated through a graded ethanol series (30%, 50%, 70%, 75%, 80%, 85%, 90%, 95%, 98%, and 100%). Samples were then dried using a critical point dryer, mounted on specimen stubs, sputter-coated with gold, and examined by scanning electron microscopy (SEM) at an accelerating voltage of 3 kV.

**Cytotoxicity assay**

Cytotoxicity was assessed using a Cell Counting Kit-8 (CCK-8) assay. HEK293T cells were maintained in DMEM supplemented with 10% FBS at 37 °C in 5% $CO_2$. Cells were seeded into 96-well plates at $1 \times 10^4$ cells per well and cultured overnight. Cells were then treated with different concentrations of lysins for 24 h. After treatment, 10 μL CCK-8 reagent was added to each well and incubated at 37 °C for 1–3 h. Absorbance at 450 nm was measured using a Synergy H1 microplate reader, with medium-only wells as the background control. Relative cell viability was calculated as:

Relative viability (%) = [(ODsample − ODnc)/(ODbuffer − ODnc)] × 100, where ODnc is the $OD_{450}$ of the negative control, ODbuffer is the $OD_{450}$ of the HEPES-treated control, and ODsample is the $OD_{450}$ of the lysin-treated sample.

**Murine periodontitis model**

All animal experiments were approved by the Animal Ethics Committee of the Wuhan Institute of Virology, Chinese Academy of Sciences (WIVA17202301). Female BALB/c mice (6-8 weeks old) were acclimated for 1 week and randomly assigned to eight groups (n = 5 per group): Healthy, Control, Minocycline, LysPd078, PlyPd06, PlyPd19, PlyPd27, and PlyPd44.

To deplete the native oral microbiota, mice received kanamycin (500 μg/mL) in drinking water for 4 days, followed by a 3-day washout period. To induce periodontitis, a sterile 5-0 silk ligature was placed around the cervical margin of the right maxillary second molar. From day 0 to day 13, all groups except the Healthy group were topically inoculated once daily around the right maxillary second molar with 100 μL of a mixture containing *P. gingivalis* W83 and SP samples (approximately $10^9$ CFU/mL, suspended in 2% carboxymethyl cellulose). From day 7 to day 13, body weight was recorded daily, and mice received topical treatment with 100 μL of 10 mM HEPES, minocycline (Periofeel Dental Ointment 2%, Showa Yakuhin Kako Co), 20 μg LysPd078, 20 μg PlyPd06, 20 μg PlyPd19, 20 μg PlyPd27, or 20 μg PlyPd44. All mice were weighed daily. On day 14, mice were euthanized, and the right maxilla, heart, liver, lungs, and kidneys were collected for further analyses.

**Micro-computed tomography analysis**

Alveolar bone loss was evaluated by high-resolution micro-computed tomography (micro-CT). Scanning was performed at 60 kV and 200 μA using a 0.5 mm Al filter, with the resolution of 4032 × 2688 and thw voxel size of 9 μm. Raw data were reconstructed using NRecon software, and three-dimensional images were generated using CTvox. In two-dimensional reconstructed images, the distance from the cementoenamel junction (CEJ) to the alveolar bone crest (ABC) of the maxillary second molar was measured using DataViewer. Bone volume fraction (BV/TV) was quantified using CTAnalyzer.

**Histological analysis**

Maxillary and calvarial samples were fixed in 4% paraformaldehyde and decalcified in 10% EDTA. Representative bone specimens were paraffin-embedded and sectioned for histological analysis. To evaluate the anti-inflammatory and tissue-protective effects of lysin treatment, sections were subjected to IL-1β immunohistochemistry, hematoxylin and eosin (HE) staining, tartrate-resistant acid phosphatase (TRAP) staining, and Masson staining. To assess systemic safety, the heart, liver, lungs, and kidneys were also collected, fixed, and subjected to H&E staining.

**Statistical analysis**

Experimental and computational data were analyzed using SPSS 26.0 and GraphPad Prism 8.0, respectively. Details of the statistical tests used for each comparison are provided in the corresponding figure legends. Data are presented as mean ± SD. Statistical significance was defined as $P < 0.05$, $P < 0.01$, $P < 0.001$, and $P < 0.0001$; ns indicates no significant difference.

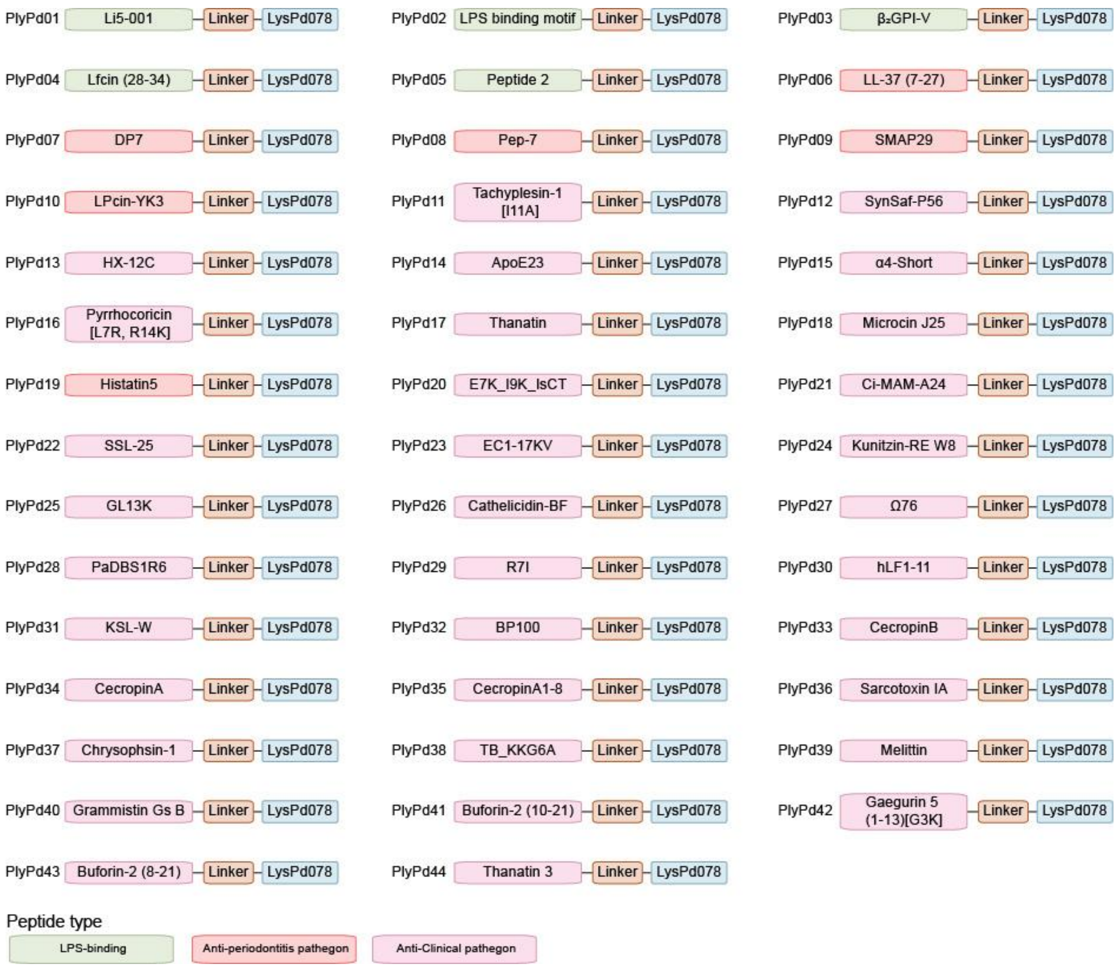


Supplementary Fig. 1 Schematic diagrams of designed PlyPds.

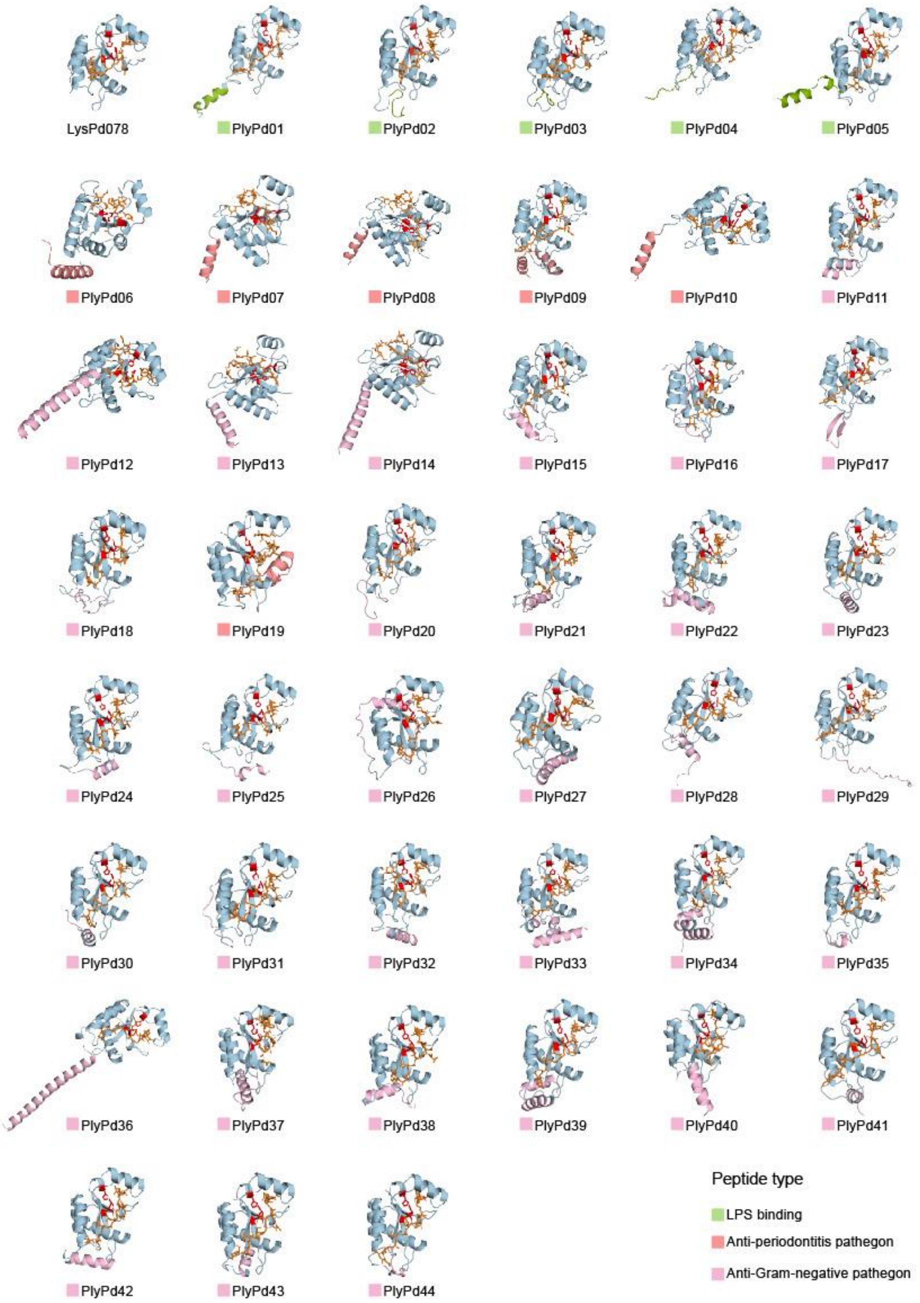


Supplementary Fig. 2 The binding conformations of LysPd078 and the designed PlyPds with PG.

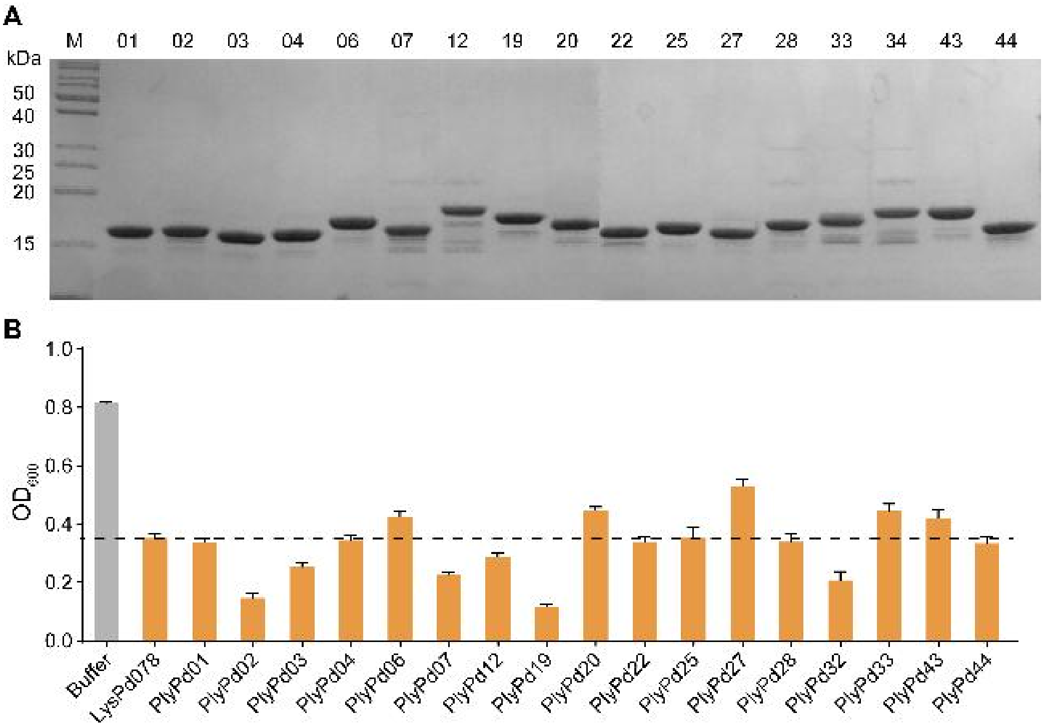


Supplementary Fig. 3 Primary screening and characterization of PlyPds. (A) 15% SDS-PAGE analysis of purified PlyPds. M: Protein molecular weight marker. (B) Degradation of P. gingivalis W83 PG by LysPd078 and selected PlyPds. The dashed line represents the average level of PG degradation by LysPd078.

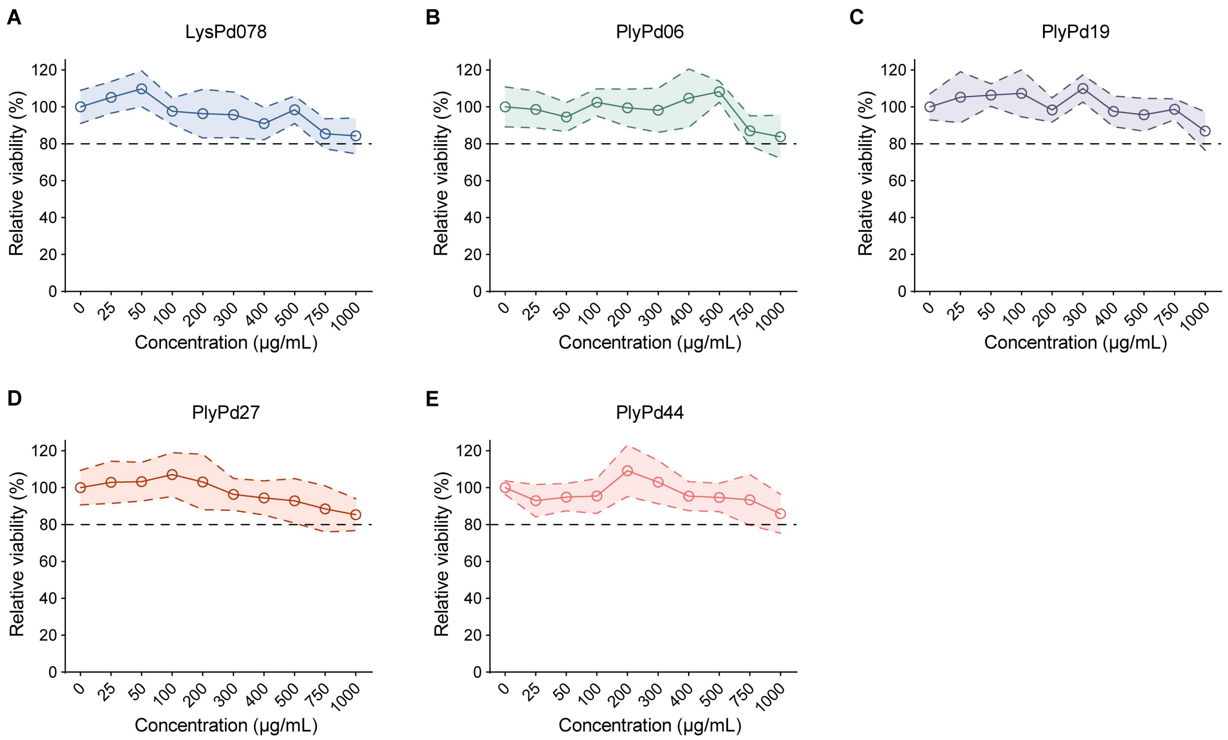


Supplementary Fig. 4 Cytotoxicity of LysPd078 (A) and the preferred PlyPds, including PlyPd06 (B), PlyPd19 (C), PlyPd27 (D), and PlyPd44 (E) on HEK293T cells. Lines and shaded regions indicate means and SD, respectively. Cell viability above 80% (dashed line) indicates no cytotoxicity.

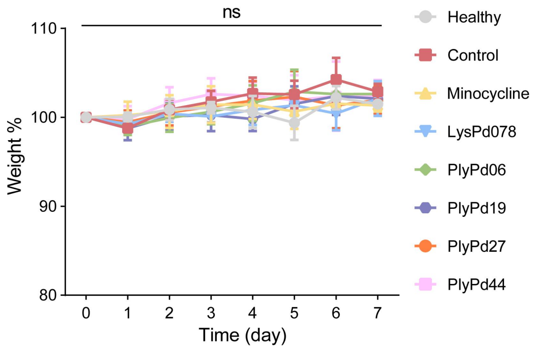


Supplementary Fig. 5 Changes in body weight among Healthy mice and those treated with different conditions. Statistical significance was calculated according to Two Way ANOVA; n = 5/group, ns, not significant. Data are presented as mean values ±

SD.
